# Impact of narrow spectrum Penicillin V on the oral and fecal resistome in a young child treated for otitis media

**DOI:** 10.1101/672196

**Authors:** Kjersti Sturød, Achal Dhariwal, Ulf R Dahle, Didrik F Vestrheim, Fernanda C Petersen

## Abstract

**Background:** Antibiotic overuse has led to a global emergence of resistant bacteria, and children are among the frequent users. Most studies with broad-spectrum antibiotics show severe impact on the resistome development of patients. Although narrow-spectrum antibiotics are believed to have less side-effects, their impact on the microbiome and resistome is mostly unknown. The aim of this study was to investigate the impact of the narrow-spectrum antibiotic phenoxymethylpenicillin (Penicillin V) on the microbiome and resistome of a child treated for acute otitis media (OM).

**Methods:** Oral and fecal samples were collected from a one-year child before (day 0) and after (day 5 and 30) receiving Penicillin V against OM. Metagenomic sequencing data was analysed to determine taxonomic profiling, using Kraken and Bracken software, and resistance profiling, using KMA in combination with the ResFinder database.

**Results:** In the oral samples, 11 antimicrobial resistance genes (ARGs), belonging to four classes, were identified at baseline. At day 5, the abundance of some ARGs was increased, some remained unchanged, while others disappeared. At day 30, most ARGs had returned to baseline levels, or lower. In the fecal samples we observed seven ARGs at baseline and five at day 5, with only one gene observed at day 5 being present at baseline. At day 30 the number of ARGs increased to 21 ARGs from 7 different classes.

**Conclusions:** Penicillin V had a remarkable impact on the fecal resistome indicating that even narrow-pectrum antibiotics may have important consequences in selecting for a more resistant microbiome.

## Introduction

Increasing prevalence of antibiotic resistant bacteria is now threatening what have been modern medicines’ security blanket for over 70 years, namely effective antibiotics ^1^. Antibiotics are among the most essential and prescribed drugs in the world, but overuse and misuse of antibiotics has led to a global emergence of resistant bacteria making bacterial infections once again difficult to treat. Antibiotic use is particularly high in children less than 4 years of age ^2,3^. The leading cause for prescriptions in these children is upper airway infections, including otitis media (OM) ^3,4^. In Scandinavia, the treatment-guidelines for OM are much more restrictive than they were some years ago ^3,5^, and doctors are encouraged to use as little antibiotics as possible in these cases. However, when antibiotics are recommended, the first choice of treatment is phenoxymethylpenicillin, also known as Penicillin V ^6,7^. Penicillin V has antibiotic activity against gram-positive bacteria, but is less active against gram negative bacteria, and it is known to be one of the narrowest spectrum antibiotics in use.

Use of broad spectrum antibiotics are associated with a reduction in gut microbiome diversity, as demonstrated in several studies ^8,9^. The impact on the microbiota in other body sites is less known, but the microbiome in saliva seems to be more resilient to antibiotics than the gut microbiome, at least in healthy volunteers exposed to broad-spectrum antibiotics ^10^. In children, the impact of loss in diversity is of particular importance, given the relevance of the microbiome on immune system maturation ^11^. While such antibiotic effects on microbial composition have been the focus of several investigations, less is known about the effects on the overall antibiotic resistance gene load in the human microbiome, known as the resistome ^12^. The resistome constitutes an important reservoir of antibiotic resistance genes that can be readily transferred to pathogens ^13^. Emerging data indicates that broad spectrum antibiotics can lead not only to a general increase in resistance genes related to the antibiotic target, but also to other groups of antibiotics ^14^. Such resistance genes can in some cases persist for long-periods, and increase upon cumulative exposure to antibiotics ^15^. For narrow-spectrum antibiotics like Penicillin V, the impact on the resistome remains largely unknown.

In the present study, we followed the effect of penicillin V on the oral and the gut resistome of a young child treated for OM. We used deep metagenomic sequencing to investigate the possible relevance of narrow spectrum antibiotics in promoting enrichment of pathogens and resistance genes in the microbiome. The results indicate that narrow spectrum antibiotics impacted both the microbiome and the resistome, especially in the gut, leading to shifts in the taxonomic profile and ARG enrichment.

## Materials and methods

### Ethics

The study was approved by the Regional Committee for Medical and Health Research Ethics – South East (“REK sør-øst”). Written, informed consent was obtained from the participating child’s parents. The participants received no compensation.

### Sample collection

Fecal and oral samples from a one-year-old child that presented with OM at the Accident and Emergency Department at Bærum hospital were collected for shotgun metagenome sequencing. The child was of Caucasian origin, had no travel history outside the Nordic Countries and had previously received antibiotic treatment at one occasion (> 1month ago). Samples were collected at 3 time points: (1) the same day as the visit to the emergency department (baseline), (2) immediately after the treatment course was completed (day 5) and (3) after 25 days post antibiotic treatment (day 30). Oral samples were collected with Copan Floqswab (Copan Diagnostics, Murrieta, CA). The swab was stored in 2 ml TE-buffer, put directly on ice and DNA was extracted within two hours of sample collection. Fecal samples were collected in sterile containers and put directly in a −20°C freezer. Fecal DNA was extracted after all samples were collected.

### DNA extraction and sequencing

DNA from the oral samples were extracted using the MasterPure Gram Positive DNA Purification Kit (Epicentre, Madison, WI). 2 ml TE-buffer containing the sample material was pelleted by centrifugation (10000 rpm for 10 minutes) and extracted by following the protocol given by the manufacturer. Fecal DNA extraction was performed using the PowerLyzer™ Powersoil^®^ DNA isolation Kit (MoBio Laboratories, Carlsbad, CA). Minor adjustments were implemented to the protocol according to the Human Microbiome Project (Core Microbiome Sampling, Protocol A, HMP Protocol #07)^16^. The amount of both oral and fecal DNA extracted was measured by NanoDrop (NanoDrop 2000c, Thermo Scientific). DNA quality was evaluated using agarose-gel electrophoresis. Metagenomic shotgun sequencing was conducted at the Norwegian Sequencing Centre (NSC) on an Illumina HiSeq 2500 platform using paired-end sequencing approach with a targeted read length of 125 bp in high-output mode.

### Quality control and preprocessing of metagenomic data

Quality of raw sequencing reads were assessed using the FastQC tool ^17^, with all samples passing the commonly applied quality criteria. Firstly, low-quality reads and adapter sequences were trimmed out using Trimmomatic ^18^. Secondly, human DNA was removed by extracting all reads mapping to an assembly of human genome (version GRch38, downloaded from NCBI GenBank) using BowTie2 ^19^. The remaining processed reads were subsequently used to characterize the microbiome and resistome in oral and fecal samples.

### Taxonomic profiling

Paired non-host reads were classified and assigned taxonomic labels using Kraken (v1) software ^20^. Reads (k-mers) were aligned against prebuilt MiniKraken DB_8GB database which encompasses complete archaeal, bacterial and viral genomes in NCBI’s RefSeq (Oct. 18, 2017). Kraken assign reads to lowest common ancestor (LCA) (genus-level rather than species) thus inaccurate species abundance estimation results, particularly in case of very similar species. To overcome this issue, the species abundance was re-estimated using Bracken ^21^. The taxonomically classified reads were visualized using Pavian and Krona charts ^22^. Further, Shannon or Chao1 diversity indexes were calculated at species-level for oral and fecal samples. The samples were rarefied to even sequencing depth (per million reads) for comparative analysis.

### Identification and Quantification of Resistance genes

The KMA program ^23^ was used in combination with ResFinder 2.1 database ^24^ to identify resistance genes in the metagenomic dataset. The ResFinder database consists of ∼2200 resistance genes which are curated from existing databases and published literature. Genes that facilitate resistance phenotypes due to point mutations were omitted, since short reads do not always distinguish between a wild-type gene and a resistant-gene variant. KMA was selected over other alignment tools (SRST2, MGmapper etc.) because of its accuracy in aligning short reads against redundant databases. Additionally, this method is able to resolve the issue of multiple read matches by evaluating and statistically testing the global alignment scores. Quality-controlled reads were mapped against ResFinder database using KMA with 80% identity threshold for resistance gene identification (Supplementary Data Sheet 1: for detailed information). To reduce the number of false-positives, only significant genes (p-value <0.05) with a coverage greater than 50% were used for further analysis. Also, the source (organism) of genes (identified template) were manually searched through their Accession ID in NCBI.

Further, the given depth/abundance of each resistance genes were normalized using RPKM (Reads Per Kilobase Million mapped reads) method for comparison across samples. Similarly, the abundance profiles were also calculated by categorizing resistance genes at antibiotics class-level. These abundance profiles were represented in the form of heatmap for both oral (Figure 3A) and fecal (Figure 3B) samples. Heatmaps were generated using Pheatmap package in R. Likewise, all the co-occurrence or correlation line plots between the ARGs and microbial taxa were created in R.

**Figure 1:**
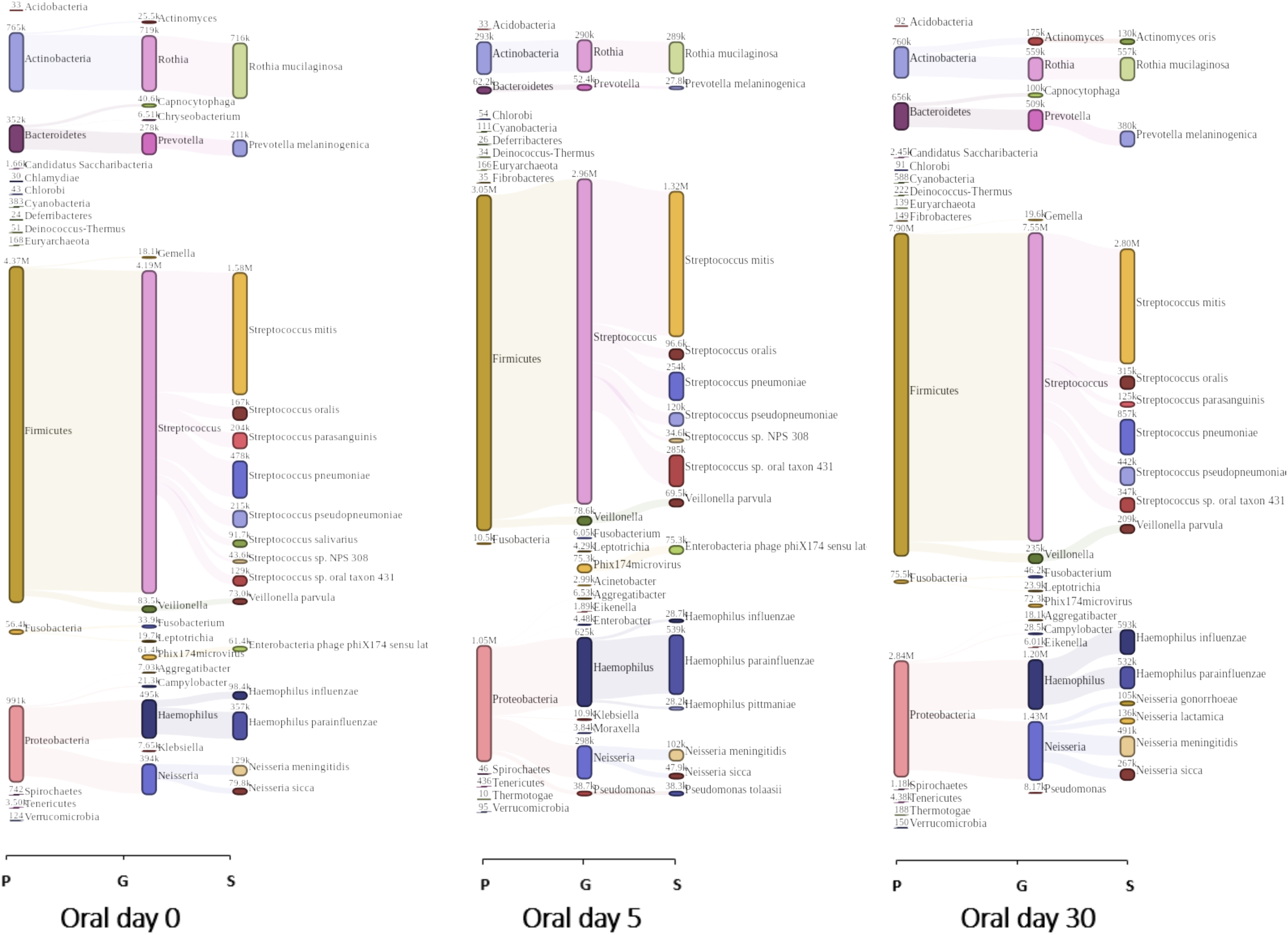
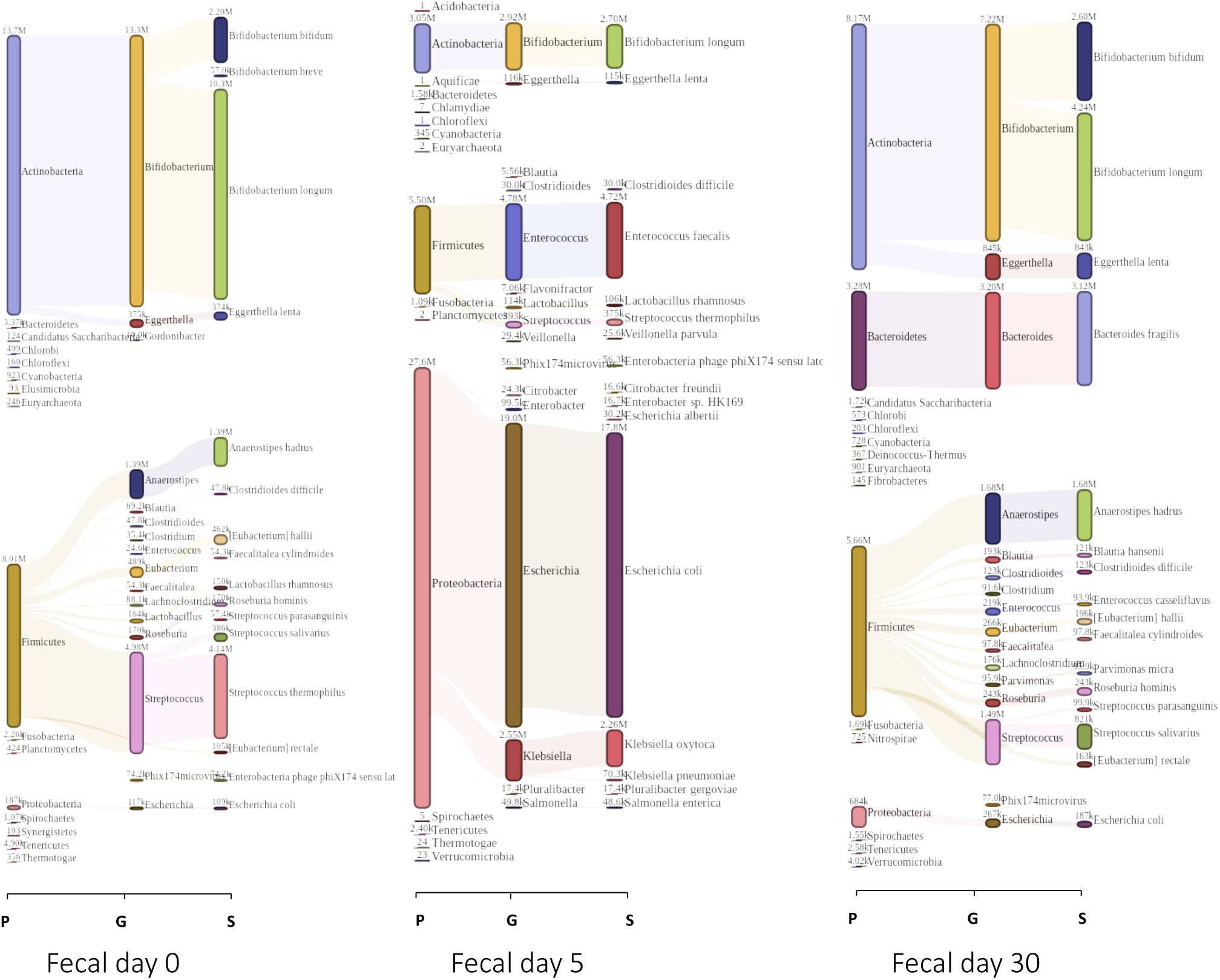
Taxonomic profile of the oral (A) and fecal (B) metagenomes classified by Kraken and visualized by Pavian. Classified reads are represented at 3 taxonomic levels (P – phylum, G – genus, S – species). The number of reads assigned to each taxon (annotated on the top of bars) is proportional to their height in Sankey diagram.

**Figure 2:**
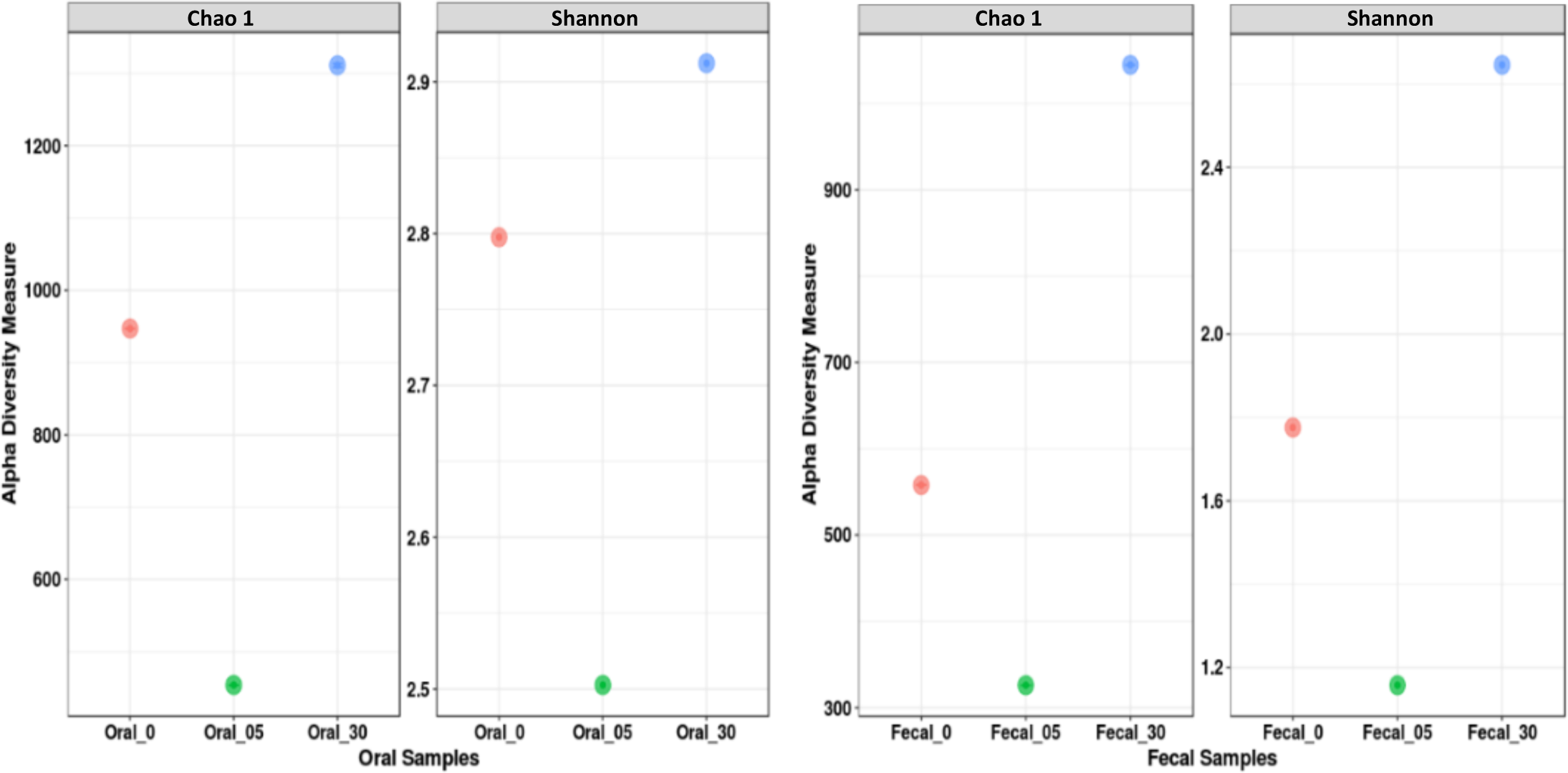
Alpha-diversity, measured by Chao1(richness) and Shannon diversity Index (evenness) is plotted for oral and fecal samples at three time points - baseline (0), immediately after treatment was completed at day 5 (05) and 25 days post antibiotic treatment (30). Both, oral and fecal samples have been rarefied to even sequencing depth for diversity estimation.

**Figure 3:**
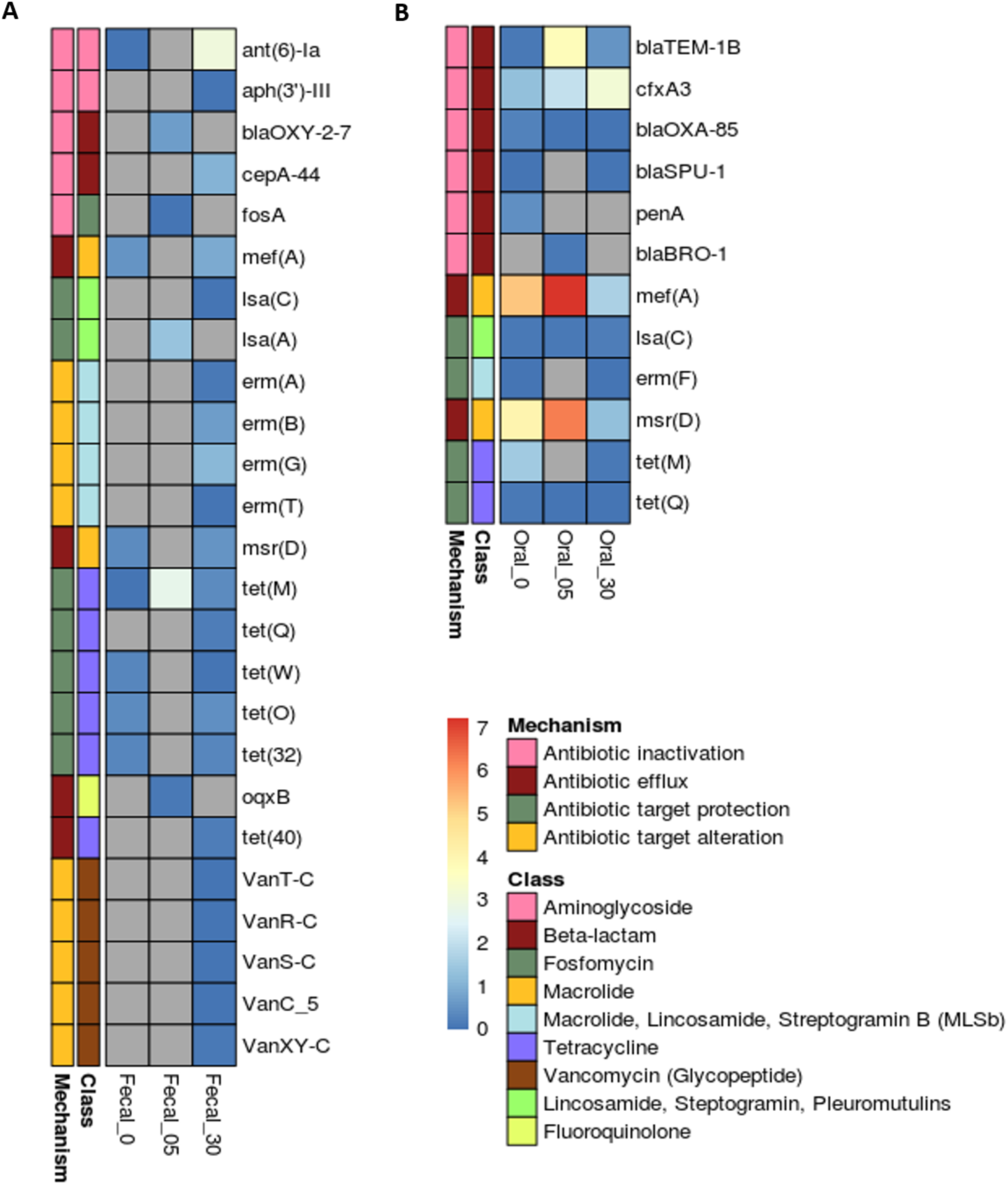
Antibiotic resistance genes identified in oral (A) and in fecal (B) samples. Heatmap showing distribution of ARGs detected within the metagenomic samples at three time points - baseline (0), immediately after the treatment course was completed at day 5 (05) and 25 days post antibiotic treatment (30). Genes are grouped by antibiotic class and mechanism of action. Colors shown in the legend (right) indicated the abundance of ARGs scaled from “blue” (minimum) to “red” (maximum). The scale 0-7 are normalized levels for the abundance of genes based on sequencing depth and gene length, as determined by RPKM. Genes which have not been detected or excluded based on cutoff criteria are colored grey.

## Results

### Sequencing results

For all 6 samples, a total of 260 million paired-reads were generated from shotgun metagenomic sequencing. The mean Phred quality score of raw reads across all samples was 37.3. Host DNA contamination was higher in oral as compared to fecal samples. On average, 66% (28M) and 8% (3M) reads were identified as human DNA in oral and fecal samples, respectively. Human DNA was filtered from the sequencing data during analysis, as described in the material and methods section above.

### Oral microbiome composition

On average, 48% reads in oral samples remained unclassified as bacteria, archaea, or viruses. The remaining taxonomically classified reads were represented by Sankey diagrams using Pavian, as shown in Figure 1A. The results suggest that the taxonomic composition of the oral metagenomes following antibiotic treatment varied slightly with respect to genus and species, but less with respect to phylum. All the oral samples were dominated by gram positive bacteria, most of them belonging to the phylum Firmicutes. The baseline (O-d0) sample contained around 67% Firmicutes, with strong dominance of the genus Streptococcus (64%). The majority of the reads were assigned to *Streptococcus mitis* (24%) at species-level. Additionally, 15%, 12% and 5% of the reads were also assigned to bacteria from the phyla Proteobacteria, Actinobacteria and Bacteroidetes, respectively. The second most abundant species in the baseline sample was *Rothia mucilaginosa* (11%) which belongs to the phylum Actinobacteria. The sample taken at day 5 (O-d5), corresponding to the day of therapy termination, shows very much the same pattern as O-d0, but with some differences. Firmicutes was still the dominating phyla, however the relative proportion of Proteobacteria was doubled compared with O-d0, and Actinobacteria, and Bacteroidetes were both reduced in their relative abundance compared with the baseline sample (O-d0). The most abundant species was still *S. mitis*; conversely, the proportion of some of the gram-negative species such as *Haemophilus parainfluenzae* increased from 5% in O-d0 to 12% in the O-d5-sample. The sample collected 25 days after the end of antibiotic exposure (O-d30) reveals a taxonomic composition which is relatively similar to O-d0 as compared to O-d5 sample. While the fraction of reads belonging to Proteobacteria were still higher as in O-d5, the relative proportion of Bacteroidetes was restored to the level shown in O-d0. Within the Proteobacteria, the highest proportion of reads were assigned to genus Neisseria and Haemophilus. Also, the relative abundance of Neisseria was found to be highest at O-d30 (12%). Furthermore, the Chao1 (richness) and Shannon (evenness) species diversity measurements decreased drastically at O-d5 compared to O-d0, and it increased again at O-d30 (Figure 2). Statistical significance of such changes could not be measured due to one child (case) in the study.

### Fecal microbiome composition

Large differences in microbiota composition were observed in the fecal samples before (F-d0), at termination (F-d5) and 25 days after (F-d30) antibiotic termination (Figure 1B). The baseline sample (F-d0) was strongly dominated by the phylum Actinobacteria (62%) and Firmicutes (36%). While at species level, large proportion of reads (47%) were assigned to *Bifidobacterium longum*. The second most abundant species was *Streptococcus thermophilus* consisting of around 19% of the total fraction of reads in the sample. However, after five days of antibiotic treatment the taxonomic profile showed a shift towards the phylum Proteobacteria completely dominating the sample with a contribution of over 76% in total reads count, compared with < 1% at baseline. A massive increase in abundance of *Escherichia coli* (49%) contributed particularly to the high proportion of Proteobacteria. In the F-d0 sample the species belonging to the Firmicutes were mainly from the genus Streptococcus or Anaerostipes, while Enterococcus (specially *Enterococcus faecalis*) was the leading genus in the Firmicutes in the F-d5-sample. In the follow-up sample from day 30 (F-d30), the relative abundance of Proteobacteria decreased to 4% as we observed in F-d0 and with Actinobacteria again dominating the sample, however, not as extensive as in the F-d0 sample. The Firmicutes were more or less back to its original composition except for *S. thermophilus*, while the phylum Bacteroidetes, with its most abundant species *Bacteroides fragilis* (17%) became a substantial part of the gut microbiome. Interestingly, *B. fragilis* was not detected in F-d0 or in F-d5. Furthermore, we observed a considerably larger portion of unmapped reads for F-d30 than for F-d0 and F-d05. For F-d05 only 10% of the reads were not mapped to a known reference compared to 58% for F-d30 and 40% for F-d0. Similar to the oral samples, the species richness and evenness in the fecal samples were reduced at F-d5 compared to F-d0 (Figure 2). Both oral and fecal samples had the highest species diversity at d30.

### Oral resistome

For oral metagenomes, we found 7 ARGs present in all samples, i.e. O-d0, O-d05 and O-d30 (Figure 3A). At baseline, a total of 11 annotated ARGs were identified at a given threshold. These genes encoded resistance to antibiotics belonging to four different classes, and used three different mechanisms of resistance. The largest group of ARGs at baseline were those that confer resistance to beta-lactams. In total, five different genes belonging to the beta-lactam group *(blaTEM-1B, cfxA3, blaOXY-85, blaSPU-1* and *penA*) were detected before antibiotic treatment started. The TEM-1 enzyme was described already in the early 1960s ^25^, and are one of the most prevalent beta-lactamase enzymes in gram negative bacteria. It is found on transferable genetic elements and is able to inactivate penicillins and narrow-spectrum cephalosporins ^26^. In addition to the beta-lactams, the efflux pump, *mef(A)*, which confers resistance to macrolides was detected at O-d0, together with *msr(D)* which is an macrolide efflux protein expressed on the same operon ^27^. The *lsa(C)* gene confer resistance to lincosamides, streptogramins and pleuromutulins, while *erm(F)* confers resistans towards the MLSB –group (macrolides, lincosamides and streptogramin B). Two genes conferring tetracycline resistance, *tet(M)* and *tet(Q)*, were also detected at O-d0. Most ARGs discovered at O-d0 were also identified at O-d5. Levels of *blaOXA-85, lsa(C)* and *tet(Q)* were similar to O-d0, however, the abundance of *blaTEM-1B, cfxA3*, mef(A) and *mrs(D)* was increased at O-d5. Of the four genes with an increased abundance after five days, different responses were observed at O-d30. *mef(A)* and *msr(D)* both decreased in abundance below their baseline level, while *cfxA3* increased even more at O-d30. *blaTEM-1B* decreased from the high abundance at O-d5, but not to the lowest levels observed at day 0. Thus, in oral samples, most of the observed changes during the antibiotic treatment returned to baseline levels at O-d30 (Figure 3A).

### Fecal resistome

From all the fecal samples, we detected in total 25 different ARGs. At baseline, 7 different ARGs were identified, conveying resistance to four different antibiotic classes (Figure 3B). *Ant(6)-Ia* encodes aminoglycoside resistance, *mef(A)* is a macrolide efflux pump, *msr(D)* is associated with resistance towards the MLSB –group and *tet(M), tet(W), tet(O)* and *tet(32)* are ribosomal protection proteins avoiding damage from tetracyclines. Only one of these genes were identified again after five days, *tet(M)*, thus, showing a shift in the fecal resistome at day 5. The abundance of *tet(M)* was increased at F-d5 compared with the baseline sample. In addition to *tet(M)*, 4 new ARGs were identified at F-d5. One beta-lactamase (*blaOXY-2*), a fosfomycin resistance gene (*fosA*), and two efflux pumps, *lsa(A)* and *oqxB*, which show resistance to macrolides and fluoroquinolones respectively. In the last sample we identified 21 ARGs in total, showing a considerably increase from the first two fecal samples. At F-d30 ARGs from seven different antibiotic classes were identified, including genes encoding resistance towards vancomycin, which were not detected at F-d0 nor at F-d5. *Tet(M)* was the only gene detected at each timepoint, with its highest abundance at F-d5. Interestingly, 14 ARGs were solely identified at F-d30, 25 days after the last antibiotic exposure. Hence, the fecal resistome showed an extensive shift in composition and increase in diversity following the antibiotic treatment.

### Relationships between the resistome and microbiome

Many of the observed ARGs, like *blaBRO-1, blaOXA-85* and *PenA* in the oral samples, and *blaOXY-2-7* and the Van-genes in the fecal samples, are found intrinsicly in a specific species, while others, like *blaTEM-1*, Tet-genes and most Erm-genes are found either intrinsicallly in several bacteria or on mobile DNA elements. Here we did a correlation analysis on resistance genes that typically can be found intrinsically in one bacteria to find potential associations between the taxa and ARG. For this, we plotted their abundances (co-occurance) across the three different timepoints in oral and fecal samples (Figure 4). In the oral samples, *blaBRO-1* which is found chromosomaly in *Moraxella catarrhalis* showed similar changes in abundance profile as *M.catarrhalis* suggesting that the increase of *blaBRO-1* on day 5 correlates with the concurrent increase of *M.catarrhalis*. The same goes for *blaOXY-2-7* and the Van-genes that are intrinsic to *Klebsiella oxytoca* and *Enterococcus casseliflavus*, respectively. *E. casseliflavus* is known to harbor intrinsic, non-transferable resistance to vancomycin due to a chromosomally encoded VanC gene cluster^28,29^. All five vancomycin resistance genes found in this study are gene variants found in this VanC cluster. *E. casseliflavus* is not as well described as *E. faecalis* and *Enterococcus faecium*, however, it can cause serious infections, and due to its high level of intrinsic resistance towards several antibiotics it is a difficult bacteria to treat^30^.

**Figure 4:**
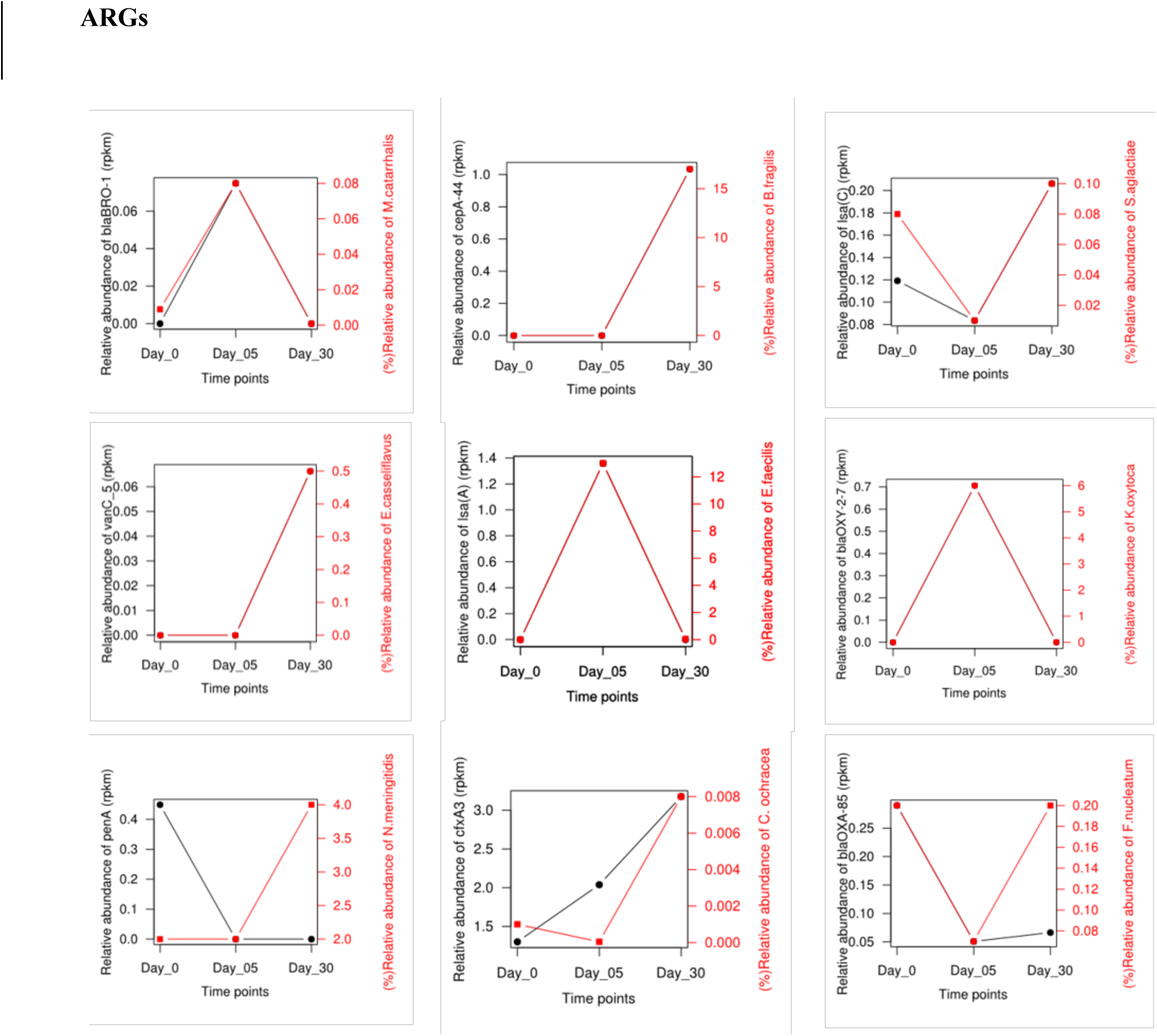
Line plots showing the abundance of ARGs (RPKM) in the oral and fecal metagenome samples (black), together with the relative abundance of selected species potentially carrying the ARGs (based on annotation by KMA) (red).

Not all genes followed the expected source of origin. The resistance genes *PenA* and *blaOXA-85* (also named *FUS-1*), which usualy are found in *Neisseria menigitidis* and *Fusobacterium nucleatum* respectively, did not follow the abundance profile of their expected host, suggesting that these genes may be intrinsic also in other bacteria, or that the genes can be found on mobile elements. For genes that are found intrinsically in several bacteria or on mobile DNA elements, it is not possible to correlate them to bacterial species with the methods applied in this study.

## Discussion

Young children are frequent users of antibiotics. In the Scandinavian countries the use of antibiotics is still lower than in many other European countries, and the low usage most likely contributes to the low prevalence of antibiotic resistance that is observered^31^. Penicillin V is recommended as first-line drug for empirical treatment of OM in Norwegian treatment guidelines^6^.

In this study, we used a sequenced based metagenomic approach to illuminate the impact Penicillin V had on the oral and fecal microbiome and resistome of a child living in Norway. Supporting previous findings^10^, our results show that there was a remarkable difference in how antibiotics affected the oral microbiota in contrast to the fecal microbiota. The oral microbiota was relatively stable towards antibiotic exposure, despite the fact that Penicillin V has an antimicrobial activity on gram-positive bacteria, which predominate in the oral cavity. Because the child had a middle ear infection, it is possible that the oral bacterial community at baseline was altered from its normal state. However, the stability of the oral samples from baseline to 25 days post antibiotic therapy indicated that the bacterial infection did not have a significant impact on the oral microbiome. Previous studies have reported the stability of the oral microbiome in response to antibiotic therapy^10^, however, previous observations have been in healthy adults, and not in young children with an infection. The current results demonstrated that the impact of Penicillin V on the oral microbiota composition in this young child was limited.

Similar to oral samples, microbial diversity in the stool samples decreased after 5 days of treatment, and after 30 days the level of diversity was higher than baseline levels. However, fecal diversity was overall lower than in oral samples, and the alterations of the taxonomic profile following antibiotic therapy were most prominent in the fecal samples, especially at day 5. In the fecal baseline sample, Bifidobacterium species predominated. In contrast, after five days of treatment, the potentially pathogenic Enterobacteriaceae were the dominating family, and the abundance of Bifidobacterium species were reduced. This is likely an undesirable outcome, since high levels of Bifidobacterium are associated with several health benefits^32^, including reduced levels of AMR in early life^33^. Conversely, Enterobacteriaceae are a group of bacteria known to harbor high levels of ARGs, and they have the highest priority by WHO in the fight against antibiotic-resistant bacteria^34^. The observed reduction of Bifidobacteriaceae, and increase in Enterobacteriaceae during Penicillin V treatment, supports previous studies investigating the effect of other beta-lactams of broader spectrum on the fecal microbiome^35,36^.

Five ARGs (*mef(A), lsa(C), msr(D), tetM* and *tet(Q)*) were observed both in the oral and the fecal microbiome. Four of these (*mef(A), msr(D), tetM* and *tet(Q)*) can be found in several bacteria on mobile elements, and are therefore of particular importance for spreadning via horizontal gene transfer^37^. The impact of Penicillin V on the resistome was greater in the fecal samples than in the oral samples. While in the oral samples, the 10 ARGs detected at O-d30 were also observed at baseline (Figure 3A), in the fecal samples, we observed 14 genes that were exclusive to d30, (Figure 3B). Interestingly, only one of the fecal ARGs identified at day 30 conferred resistance towards beta-lactam antibiotics, which was used to treat the patient. ARGs conferring resistance towards beta-lactam antibiotics was also observed in oral samples, but these were not enriched following antibiotic therapy. The extensive impact on the fecal resistome at day 30 compared with the modest effects on the oral sample might be explained by a larger taxonomic shift in the fecal samples than in the oral samples with new taxa, like *E. casseliflavus* and *B. fragilis*, contributing to new genes in the resistome. However, the large difference in the resistome of the fecal samples may also be due to higher antibiotic selection pressure and horizontal gene transfer within the already existing microbiota^38^.

Although definitive conclusions require a larger and ideally randomized study, this case report indicates that even narrow spectrum antibiotics may have a significant effect on the fecal microbiome and resistome, thus warranting further investigations. Overall, acute and recurrent OM are among the most predominant conditions associated with antibitotic prescription, and thus studies on these patients should be encouraged.

## Acknowledgment

We thank Germar Schneider and the Accident & emergency department at Bærum Hospital for contributions in sample collection. This work was partly funded by the Dental Faculty, University of Oslo and the Norwegian Institute of Public Health. AD is supported by a PhD grant from the Research Council of Norway (grant number 421258) The sequencing service was provided by the Norwegian Sequencing Centre (www.sequencing.uio.no).

## References

1. Davies SC. Annual Report of the Chief Medical Officer, Infections and the rise of antimicrobial resistance London: Department of Health (2013)2011.

2. Holstiege J, Schink T, Molokhia M, et al. Systemic antibiotic prescribing to paediatric outpatients in 5 European countries: a population-based cohort study. BMC pediatrics. 2014;14:174.

3. Størdal K, Mårild K, Blix HS. [Use of antibiotics in children during the period 2005 - 16]. Tidsskr Nor Legeforen. 2017;137(18).

4. Hersh AL, Shapiro DJ, Pavia AT, Shah SS. Antibiotic prescribing in ambulatory pediatrics in the United States. Pediatrics. 2011;128(6):1053–1061.

5. Treatment for acute inflammation of the middle ear in children. 2000; http://strama.se/wp-content/uploads/2016/04/Konsensut_ora_eng.pdf. Accessed 15.03, 2019.

6. Akutt Otitis Media. Nasjonal faglig retningslinje for antibiotikabruk i primærhelsetjenesten http://www.antibiotikaiallmennpraksis.no/index.php?action=showtopic&topic=VMpmsqDE. Accessed 14.03, 2019.

7. Solen MG, Hermansson A. Otitis media-from self-healing to life-threatening. New therapeutic guidelines. Lakartidningen. 2011;108(19):1049–1052.

8. Dethlefsen L, Huse S, Sogin ML, Relman DA. The pervasive effects of an antibiotic on the human gut microbiota, as revealed by deep 16S rRNA sequencing. PLoS biology. 2008;6(11):e280.

9. Panda S, El khader I, Casellas F, et al. Short-term effect of antibiotics on human gut microbiota. PloS one. 2014;9(4):e95476.

10. Zaura E, Brandt BW, Teixeira de Mattos MJ, et al. Same Exposure but Two Radically Different Responses to Antibiotics: Resilience of the Salivary Microbiome versus Long-Term Microbial Shifts in Feces. mBio. 2015;6(6):e01693–01615.

11. Round JL, Mazmanian SK. The gut microbiota shapes intestinal immune responses during health and disease. Nature reviews Immunology. 2009;9(5):313–323.

12. Wright GD. The antibiotic resistome: the nexus of chemical and genetic diversity. Nature reviews Microbiology. 2007;5(3):175–186.

13. Penders J, Stobberingh EE, Savelkoul PH, Wolffs PF. The human microbiome as a reservoir of antimicrobial resistance. Frontiers in microbiology. 2013;4:87.

14. Willmann M, El-Hadidi M, Huson DH, et al. Antibiotic Selection Pressure Determination through Sequence-Based Metagenomics. Antimicrobial agents and chemotherapy. 2015;59(12):7335–7345.

15. Jernberg C, Löfmark S, Edlund C, Jansson JK. Long-term ecological impacts of antibiotic administration on the human intestinal microbiota. The ISME journal. 2007;1(1):56–66.

16. Core Microbiome Sampling Protocol AHMP Protocol # 07-001. Manual of Procedures 2010; https://hmpdacc.org/hmp/doc/HMP_MOP_Version12_0_072910.pdf. Accessed 15.03, 2019.

17. Andrews S. FastQC: a quality control tool for high throughput sequence data. 2010.

18. Bolger AM, Lohse M, Usadel B. Trimmomatic: a flexible trimmer for Illumina sequence data. Bioinformatics (Oxford, England). 2014;30(15):2114–2120.

19. Langmead B, Salzberg SL. Fast gapped-read alignment with Bowtie 2. Nature methods. 2012;9(4):357–359.

20. Wood DE, Salzberg SL. Kraken: ultrafast metagenomic sequence classification using exact alignments. Genome biology. 2014;15(3):R46.

21. Lu J, Breitwieser FP, Thielen P, Salzberg SL. Bracken: estimating species abundance in metagenomics data. PeerJ Computer Science. 2017;3:e104.

22. Ondov BD, Bergman NH, Phillippy AM. Interactive metagenomic visualization in a Web browser. BMC bioinformatics. 2011;12(1):385.

23. Clausen PT, Aarestrup FM, Lund O. Rapid and precise alignment of raw reads against redundant databases with KMA. BMC bioinformatics. 2018;19(1):307.

24. Zankari E, Hasman H, Cosentino S, et al. Identification of acquired antimicrobial resistance genes. Journal of antimicrobial chemotherapy. 2012;67(11):2640–2644.

25. Datta N, Kontomichalou P. Penicillinase synthesis controlled by infectious R factors in Enterobacteriaceae. Nature. 1965;208(5007):239–241.

26. Salverda ML, De Visser JA, Barlow M. Natural evolution of TEM-1 beta-lactamase: experimental reconstruction and clinical relevance. FEMS microbiology reviews. 2010;34(6):1015–1036.

27. Ambrose KD, Nisbet R, Stephens DS. Macrolide efflux in *Streptococcus pneumoniae* is mediated by a dual efflux pump (mel and mef) and is erythromycin inducible. Antimicrobial agents and chemotherapy. 2005;49(10):4203–4209.

28. Hegstad K, Mikalsen T, Coque TM, Werner G, Sundsfjord A. Mobile genetic elements and their contribution to the emergence of antimicrobial resistant *Enterococcus faecalis* and *Enterococcus faecium*. Clinical microbiology and infection: the official publication of the European Society of Clinical Microbiology and Infectious Diseases. 2010;16(6):541–554.

29. Vincent S, Minkler P, Bincziewski B, Etter L, Shlaes DM. Vancomycin resistance in *Enterococcus gallinarum*. Antimicrobial agents and chemotherapy. 1992;36(7):1392–1399.

30. Monticelli J, Knezevich A, Luzzati R, Di Bella S. Clinical management of non-faecium non-faecalis vancomycin-resistant enterococci infection. Focus on *Enterococcus gallinarum* and *Enterococcus casseliflavus/flavescens*. Journal of infection and chemotherapy: official journal of the Japan Society of Chemotherapy. 2018;24(4):237–246.

31. Goossens H, Ferech M, Vander Stichele R, Elseviers M, Group EP. Outpatient antibiotic use in Europe and association with resistance: a cross-national database study. The Lancet. 2005;365(9459):579–587.

32. Hidalgo-Cantabrana C, Delgado S, Ruiz L, Ruas-Madiedo P, Sanchez B, Margolles A. Bifidobacteria and Their Health-Promoting Effects. Microbiology spectrum. 2017;5(3).

33. Taft DH, Liu J, Maldonado-Gomez MX, et al. Bifidobacterial Dominance of the Gut in Early Life and Acquisition of Antimicrobial Resistance. mSphere. 2018;3(5).

34. Global Priority List of Antibiotic-Resistant Bacteria to Guide Research, Discovery, and Development of New Antibiotics. World Health Organization; 2017.

35. Tanaka S, Kobayashi T, Songjinda P, et al. Influence of antibiotic exposure in the early postnatal period on the development of intestinal microbiota. FEMS immunology and medical microbiology. 2009;56(1):80–87.

36. Arboleya S, Sanchez B, Milani C, et al. Intestinal microbiota development in preterm neonates and effect of perinatal antibiotics. The Journal of pediatrics. 2015;166(3):538–544.

37. Partridge SR, Kwong SM, Firth N, Jensen SO. Mobile Genetic Elements Associated with Antimicrobial Resistance. Clinical microbiology reviews. 2018;31(4).

38. Davies J, Davies D. Origins and evolution of antibiotic resistance. Microbiology and molecular biology reviews: MMBR. 2010;74(3):417–433.

